# A glimpse at deep brain stimulation mechanisms using subthalamic nucleus optogenetic manipulations

**DOI:** 10.1101/450767

**Authors:** Alix Tiran-Cappello, Yann Pelloux, Cécile Brocard, Mickaël Degoulet, Christelle Baunez

**Affiliations:** Institut de Neurosciences de la Timone, UMR7289 CNRS & Aix-Marseille Université, 27 Boulevard Jean Moulin - 13005 Marseille, France

**Keywords:** basal ganglia, high frequency stimulation, food, cocaine, motivation

## Abstract

Although deep brain stimulation (DBS) is now a widely used therapeutic strategy, its precise mechanism remains largely unclear. Since this approach is progressively extended to treat non-motor disorders such as depression, obsessive-compulsive disorders, the comprehension of its effects on motivated behaviors appears of the upmost importance for a possible application for addiction. In intact rats, we used inhibition and high frequency optogenetic activation of subthalamic nucleus (STN) neurons to test whether or not we could reproduce the effects of electric deep brain stimulation on rats’ motivation for sweet food and cocaine. Rats’ motivation was assessed using fixed-ratio 5 and progressive ratio schedules of reinforcement for both rewards and illumination was applied during behavioral testing. Efficiency of optogenetic manipulations has been validated using in-vitro electrophysiological recordings. Optogenetic inhibition of STN increased motivation for food and reduced motivation for cocaine. In contrast, optogenetic high frequency stimulation reduced the motivation for food without impacting motivation for cocaine. Optical inhibition mimics the effect of electric deep brain stimulation on food and cocaine motivation, confirming that the effects observed under electric DBS result from a specific inactivation of the STN. In contrast, optogenetic high frequency stimulation induces opposite effects to those of electric one, suggesting a stimulation of the STN that only seems to affect food motivation.

## Introduction

The subthalamic nucleus (STN) is a small homogenous glutamatergic structure of the basal ganglia. Historically known as a relay for motor information, STN is a well-established target for deep brain stimulation (DBS) treatment for motor symptoms of Parkinson’s disease (PD) (Krack et al., 2010). STN functions have also been extended to decision-making, behavioral inhibition, reward encoding and control of motivation (Baunez et al., 2011; Hamani et al., 2017).

In rodents, the exploration of STN motivational functions has brought up interesting findings, such as the dissociative effect showing that both STN lesion and DBS increase the motivation for sweet food, while decreasing the motivation for cocaine (Baunez et al., 2005; Rouaud et al., 2010). Similarly STN pharmacological inactivation is able to reduce cocaine reinstatement (Bentzley and Aston-Jones, 2017). Finally, STN lesion and DBS have since been shown to prevent cocaine escalation and reescalation of cocaine or heroin intake (Pelloux et al., 2018; Wade et al., 2017).

In human, manipulations of the STN by DBS in PD patients have confirmed its involvement in motivated behavior such as hypersexuality (Doshi and Bhargava, 2008; Romito et al., 2002), or reduced dopaminergic dysregulation traits (Eusebio et al., 2013). STN DBS has progressively become a therapeutic strategy for the treatment of obsessive compulsive disorder (Mallet et al., 2008, 2002) and its potential to treat drug abuse disorder has been suggested (Pelloux and Baunez, 2013) and is being investigated. Despite its extensive use in these pathologies, the exact mechanism of STN DBS remains a matter of debate. For instance while being functionally similar to an inactivation in many cases, electric STN DBS locally inhibits STN cell bodies (Benazzouz et al., 2000; Beurrier et al., 2001; Meissner et al., 2005; Tai et al., 2003; Welter et al., 2004), while increasing activity of STN output structures in some studies (Hashimoto et al., 2003; Maurice et al., 2003; Shehab et al., 2014), but has also been reported to inhibit the neurons of the substantia nigra reticulata (Benazzouz et al., 2000; Tai et al., 2003). It also impacts the activity of afferent structures and namely alters cortical activity through antidromic activation of the hyperdirect pathway (Li et al., 2012; McIntyre et al., 2004).

More recently, optogenetic studies have been developed and could allow a more specific way to alter the functioning of STN, while sparing the passing fibers and neighboring structures. Using this technique, it was shown that STN DBS effects in rodent PD models are highly dependent on the hyperdirect pathway (Gradinaru et al., 2009; Sanders and Jaeger, 2016). It was later demonstrated that optogenetic inhibition of the STN was sufficient to alleviate PD symptoms in rodents (Yoon et al., 2016, 2014), thus confirming historical pharmacological inactivation and lesion studies that had shown an improvement of motor functions in PD models after STN inhibition (Baunez et al., 1995; Bergman et al., 1990; Levy et al., 2001). To date, optogenetic experiments have been used to dissect circuits involved in the motor effects of STN DBS, however motivational aspects remain largely unexplored, even though a recent study used optogenetic control to demonstrate the causal role of the STN in pausing/interrupting initiated behavior (Fife et al., 2017). In the present work we used STN optogenetic modulation to compare its effects to those previously described of electric DBS on motivated behavior in rats (Rouaud et al., 2010) and then discuss the mechanisms involved. Thus the results presented here underline the specific contribution of STN neurons to DBS mechanisms.

## Results

### Motivation for sweet food

It has been previously shown that STN electric DBS increases the motivation for sweet food, while decreasing the motivation for cocaine (Rouaud et al., 2010). To test whether STN optogenetic inhibition could affect food-motivated behavior; we injected rats bilaterally with either ARCHT3.0 (inhibition, n=9) or EYFP (Control, n=7) expressed under the control of promoter CamKIIa (Fig. 1A) and subjected to fixed ratio 5 (FR5) and progressive ratio (PR) schedules of reinforcement. Due to relatively long experimental sessions, the light was applied intermittently by 5min of laser modulation in alternation with 5 min OFF periods to reduce the risks of tissue damage (Fig. 1B). Under FR5 schedule of reinforcement, STN inactivation applied during eight consecutive sessions did not significantly change the number of food pellets obtained by the rats, although a trend towards an increase was observable at the end of the experiment and during the OFF period (Fig. 1C). Under PR schedule of reinforcement, the laser inducing inhibition transiently increased the motivation to work for sweet food reward in comparison with the level of motivation exhibited by the EYFP control group as shown by both the number of rewards obtained and the breakpoint reached (Fig. 1D). At the end of the OFF period, we performed an extra day of testing during which the inhibition induced by the laser modulation increased the breakpoint when compared to that of the EYFP control group (Fig. 1D). This also confirmed that the opsins were still functional at this stage of the experiment. Finally a detailed analysis over the first day of laser illumination and the laser challenged revealed an increased motivation for food in the inhibition group compared to the control group (Fig. 1E).

**Fig. 1:**
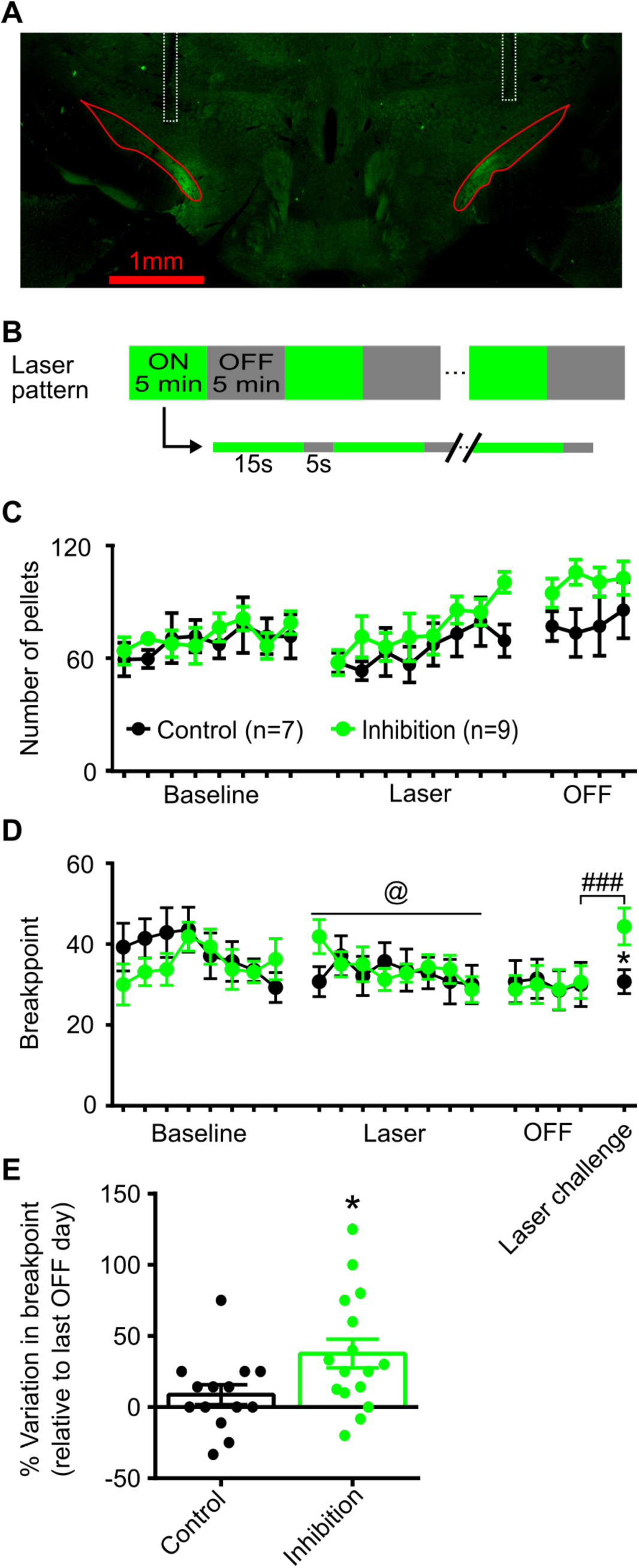
STN optogenetic inactivation increases the motivation of rats to work for sweet food. **A.** Representative image showing the fluorescence in the STN (red lines) for an animal expressing ARCHT. Dotted rectangles indicate the position of the optic fibers. **B.** Schematic representation of laser pattern used for STN inhibition, which consisted in an alternation of 5min ON (15s illumination followed by 5s of obscurity) and OFF bins, and during ON bins laser was applied using a repeating pattern of 15s illumination followed by 5s of obscurity. **C.** Under FR5 schedule of reinforcement, laser inhibition of the STN applied during 8 consecutive sessions, did not significantly alter the number of pellets obtained by the rats (Mixed ANOVA, F sessions x subjects (19, 247) = 1.550; P = 0.0699). **D.** Under PR schedule of reinforcement, laser inhibition transiently increased the breakpoint (i.e. last ratio completed) of rat working for sugar pellets (Mixed ANOVA, F sessions x subjects (7, 91) = 2.574, P = 0.0182, @ interaction effect). One day after the OFF period, another day of laser inhibition (laser challenge) was performed, which resulted in an increased breakpoint in the rats subjected to optogenetic inhibition (Mixed ANOVA, F sessions x subjects (1, 13) = 8.174, P = 0.0134, Fisher’s post-hoc test: * P <0.05, vs control, ### P < 0.001, vs OFF). **E.** Percentage of changes in breakpoint when the laser illumination was turned ON, relative to previous day without laser modulation. Data correspond to the baseline-laser transition and the laser challenge (Mann-Whitney test, P = 0.0324)

We then tested the effects of high frequency optogenetic stimulation of the STN on food-motivated behavior, by injecting the fast kinetic channelrhodopsin CHETA-TC (n=7) or the EYFP (n=5) bilaterally in the STN (Fig. 2A). The light stimulation was applied at 130Hz with a pulse-width of 2ms to mimic the electric DBS parameters in terms of frequency and using the same intermittent pattern described for inhibition experiment (Fig. 2B). In the FR5 procedure, high frequency optogenetic stimulation reduced the number of pellets obtained by the rats but only when the power output was changed from 5mW to 10mW (Fig. 2C & 2D). In the PR schedule of reinforcement, the 130Hz laser stimulation reduced the breakpoint of the CHETA-TC group compared to that reached by the EYFP control group (Fig. 2E). After a few consecutive sessions, the performance of the stimulated animals resumed to that of the control group. Interestingly when the stimulation was stopped we observed transient increase in the breakpoint of the animals that had been previously stimulated, thus indicating the presence of a rebound effect (Fig. 2E and 2F). We then tested various parameters to search for the optimal efficacy of the light stimulation. Variations of the frequencies of stimulation revealed that high frequencies (130Hz) are more efficient at reducing food-related motivation, in contrast with lower frequencies that did not modify the behavior (Fig. 2G). Finally, we tested the influence of the light pulse duration at 100Hz, by increasing the duration from 2ms to 3ms. This last manipulation further reduced motivation for sweet food (Fig. 2H).

**Fig. 2:**
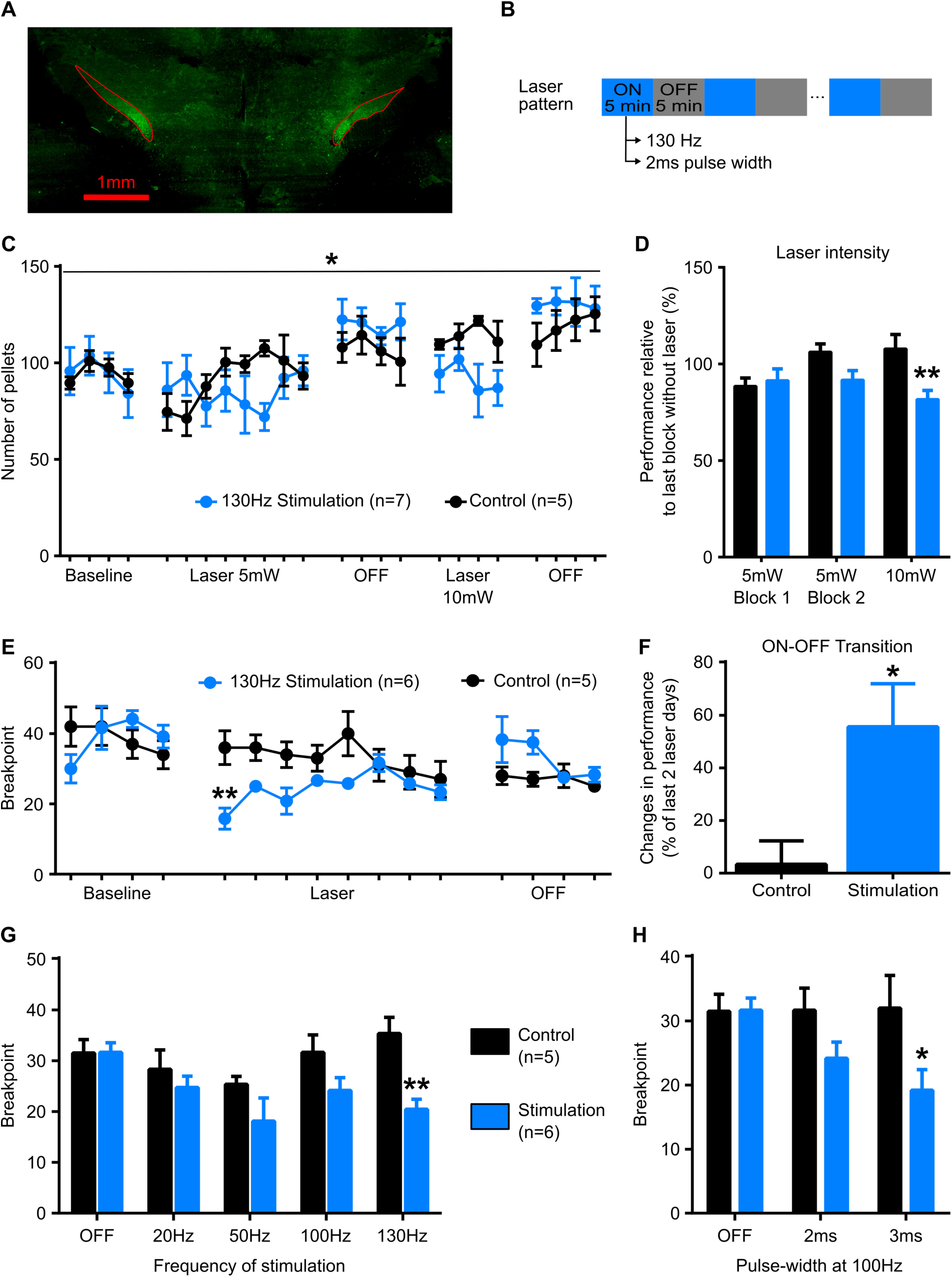
STN optogenetic stimulation at 130Hz reduces motivation for sweet food. **A.** Example of fluorescence observed in the STN (red lines) in an animal transfected with CHETA-TC. **B.** Schematic representation of laser pattern used for STN stimulation, which consisted in an alternation of 5min ON and OFF bins, during the former, the laser was applied at 130Hz without interruptions. **C.** In the FR5 task, the influence of 130Hz STN optogenetic stimulation on the number of pellets obtained by the rats was assessed over multiple sessions, application of laser stimulation with an intensity of 5mW and 10mW reduced the food intake. (Mixed ANOVA, F sessions x subjects (23, 230) = 1.794, P = 0.0168, * interaction effect). **D.** Comparison of the influence of laser intensity on rat performance calculated on 4-day blocks relative to the last block with no laser stimulation (Mixed ANOVA, F intensity _x_ subjects (2, 20) = 4.610, P = 0.0226; Sidak’s post-hoc test vs. control group: **, P < 0.01). **E.** Under PR schedule of reinforcement, the break point of stimulation group was reduced by the 130Hz optogenetic stimulation (Mixed ANOVA, Interaction F sessions x subjects (17, 153) = 2.736, P = 0.0006; Sidak’s post-hoc test vs. control group: **, P < 0.01). **F.** The performances of the 2 first OFF sessions were normalized using the last 2 sessions under optogenetic stimulation: after 8 consecutive sessions of laser stimulation, animals exhibited a transient increase in breakpoint when the laser was turned OFF (Mann Whitney test, P = 0.0476). **G.** Efficiency of laser stimulation was assessed at various frequencies revealing higher impact for high frequencies. Data were averaged over 3-day blocks (Mixed ANOVA, F frequency x subjects (4, 36) = 3.001, P = 0.0310; Sidak’s post-hoc test vs. control group: **, P < 0.01). **H.** The influence of the pulse duration on the breakpoint was assessed at 100Hz. 3ms pulses were more effective than 2ms pulses to impact food motivation in the PR task, data were averaged over 3-day blocks (Mixed ANOVA, F _pulse_ duration _x subjects_ (2, 18) = 4.670, P = 0.0232; Sidak’s post-hoc test vs. control group: *, P < 0.05).

Since the STN is primarily known for its motor functions, we performed a locomotion experiment to assess the effect of optogenetic modulation on such behavior. Laser beam activation increased the locomotor activity of the ‘inhibition’ group while it remained unaffected in the ‘stimulation’ group (Fig. S1A).

### Motivation for cocaine

Because STN DBS reduces motivation for cocaine (Rouaud et al., 2010), we trained animals of control (n=5), inhibition (n=5) and stimulation (n=5) groups to nose-poke for cocaine under FR5 schedule of reinforcement. We applied the same intermittent laser pattern with 5min ON and OFF bins described previously during the food self-administration experiment. Optogenetic inhibition of the STN reduced the cocaine intake in ARCHT animals under FR5 schedule of reinforcement (Fig. 3A). Normalizing the rat intake relative to the last day of baseline further highlighted this decrease in cocaine intake (Fig. 3B & C). In contrast, the 130Hz stimulation at 10mW did not affect the cocaine intake (Fig. 3E-G). As optogenetic inhibition can impact locomotion, we calculated the percentage of active nose-pokes performed during laser bins, and none of the stimulation or inhibition groups displayed an altered pattern compared to the control group, thus ruling out the risk of nonspecific effects related to motor activity (Fig. 3D & H). We also tested possible interactions of cocaine with laser stimulation on locomotor activity, and none of the three groups; control, inhibition and stimulation displayed any changes following the injection of a 5, 7.5 or 10mg/kg dose of cocaine (Fig. S1B-D).

**Fig. 3:**
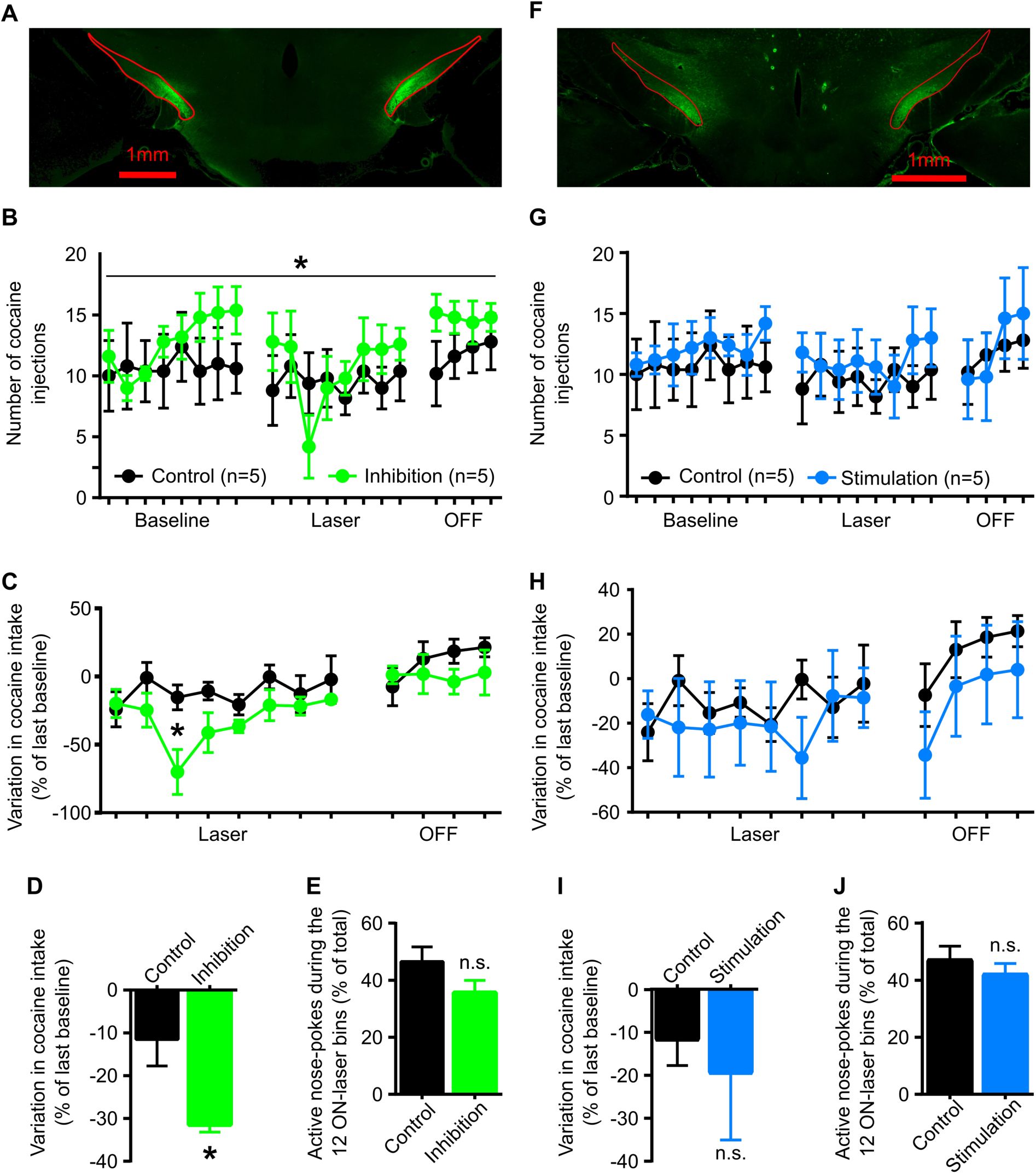
STN optogenetic inhibition but not stimulation reduces cocaine intake in FR5 task. **A.** Example of STN fluorescence in the STN (red lines) of an animal expressing inhibitory opsins ARCHT. **B.** Laser inhibition reduced the number of cocaine injections obtained by the rats in the FR5 (Mixed ANOVA, F sessions _x_ subjects (19, 152) = 1.958, P = 0.0137). **C.** Normalization using the performance during the last day of baseline further highlighted this decrease (Mixed ANOVA, F _Group_ (1, 8) = 5.772, P = 0.0430; Sidak post-hoc test: *, P < 0.05). **D.** Averaged performance across the eight sessions of laser inhibition revealed a lower level of intake compared to controls (Mann Whitney test, P = 0.0317, ###, P < 0.001 for inhibition group). **E.** The percentage of active nose-poke when the laser stimulation was turned ON (i.e. during laser bins) was calculated and was not different between stimulation and control group (Mann Whitney test, P = 0.2222). **F.** Typical image of fluorescence in the STN (red lines) of an animal expressing excitatory opsins CHETA-TC. **G.** 130Hz laser stimulation did not changed the number of cocaine injection taken by the stimulated animals compared to the control group (Mixed ANOVA, F sessions _x_ subjects (19, 152) = 0.5815; P = 0.9151). **H.** Normalization to the last day of baseline revealed a similar level of performance when controlling for inter‐ individual variability (Mixed ANOVA, F sessions x subjects (11, 88) = 0.7318, P = 0.7053). **I.** Average variation of performance over the 8 laser sessions did not emphasize any effect of 130Hz laser stimulation (Mann Whitney test, P = 0.8889). **J.** The percentage of active nose-pokes when laser stimulation was turned ON did not differ between ‘stimulation’ and ‘control’ groups (Mann Whitney test, P = 0.9999).

Under a PR schedule of reinforcement, laser inhibition of STN immediately reduced the number of injections obtained by the inhibition group and the maximum ratio reached by the rats, thus indicating a lower level of motivation for the cocaine (Fig. 4A & D). This effect persisted for a few days (Fig. 4B) and resulted in a global lower level of intake over the 8 days of laser modulation (Fig4. C). High frequency optogenetic stimulation at 130 Hz, unlike electrical 130Hz stimulation, had no effect on performance (Fig. 4E-D). Finally, we subjected the rats to acute 20Hz optogenetic stimulation, which had no impact on the motivation to work for cocaine either (Fig. S3). As during FR5 conditions, laser illumination did not alter the pattern of nose pokes in the PR task (Fig. S4).

**Fig. 4:**
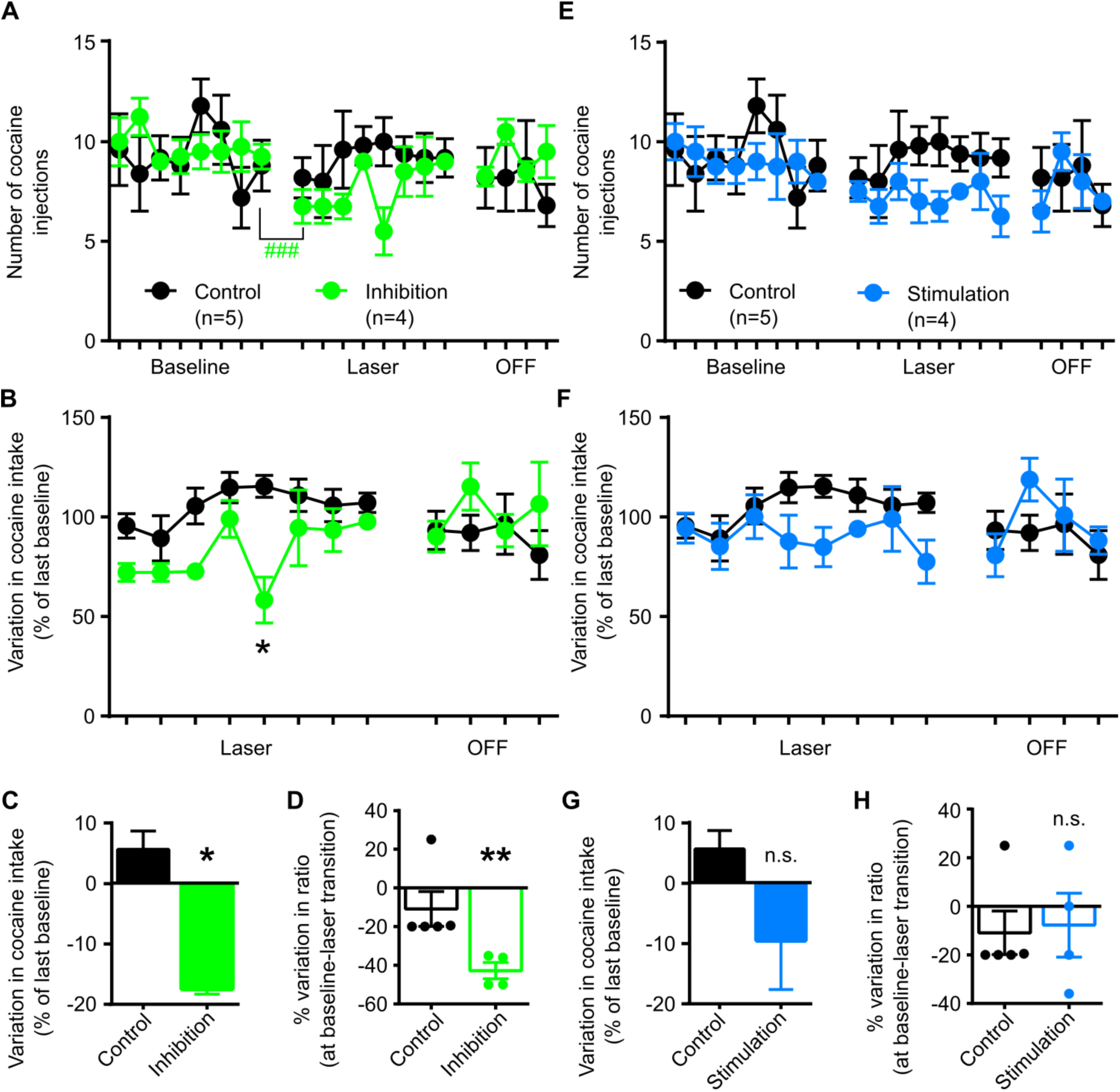
STN optogenetic inhibition reduces cocaine intake in PR task and optogenetic stimulation produces a moderate trend toward a decrease. **A.** Laser inhibition reduced the number of cocaine injections obtained by the rats in the PR task (Mixed ANOVA, F sessions x subjects (19, 133) = 1.972, P = 0.0138, ###, P < 0.0001 for inhibition group). **B.** Normalization of data by the last day of baseline allowed controlling for inter-individual variability and thus highlighted changes in cocaine intake during laser inhibition (Mixed ANOVA, F sessions x subjects (11, 77) = 2.144, P = 0.0265; Sidak post-hoc test: *, P < 0.05, vs control). **C.** Average variations in performance over the 8 laser sessions of inhibition further accentuated this decrease (Mann Whitney test, P = 0.0159). **D.** Variations in breakpoint (ratio) at baseline-laser transition for inhibition and control groups (Mann Whitney test, P = 0.0079). **E.** 130Hz laser stimulation did not change cocaine intake of stimulated animals compared to the control group (Mixed ANOVA, F sessions _x_ subjects (19, 133) = 1.185, P = 0.2792). **F.** Normalization to the last day of baseline did not highlight further variations in cocaine intake (Mixed ANOVA, F sessions _x_ subjects (11, 77) = 1.620, P = 0.1097). **G.** Average performance over laser sessions revealed a similar level of intake (Mann Whitney test, P = 0.2540). **H.** Variations in breakpoint (ratio) at baseline-laser transition for stimulation and control groups (Mann Whitney test, P = 0.9999).

### Response of STN neurons to optogenetic modulation

Ultimately, to confirm the effect of our optogenetic manipulations, we performed patch-clamp recordings on 19 adult rat brains infected with AAV virus to induce the expression of the opsins in the STN. We then confirmed that the presence of opsins did not deeply change the properties of the cells, as resting membrane potential, cell capacitance, membrane resistance, access resistance and firing rates were equivalent between stimulation, inhibition and control groups (Fig. S5A-E). To confirm the efficacy of the optogenetic modulation, we first applied discrete light pulses on STN neurons. Longer pulse-width duration and higher light intensity elicited higher inhibitory currents in ARCHT3.0 positive cells (Fig. S6A-C) and elicited higher depolarizing currents in CHETA-TC positive cells (Fig. S6D-F).

**Fig. 5:**
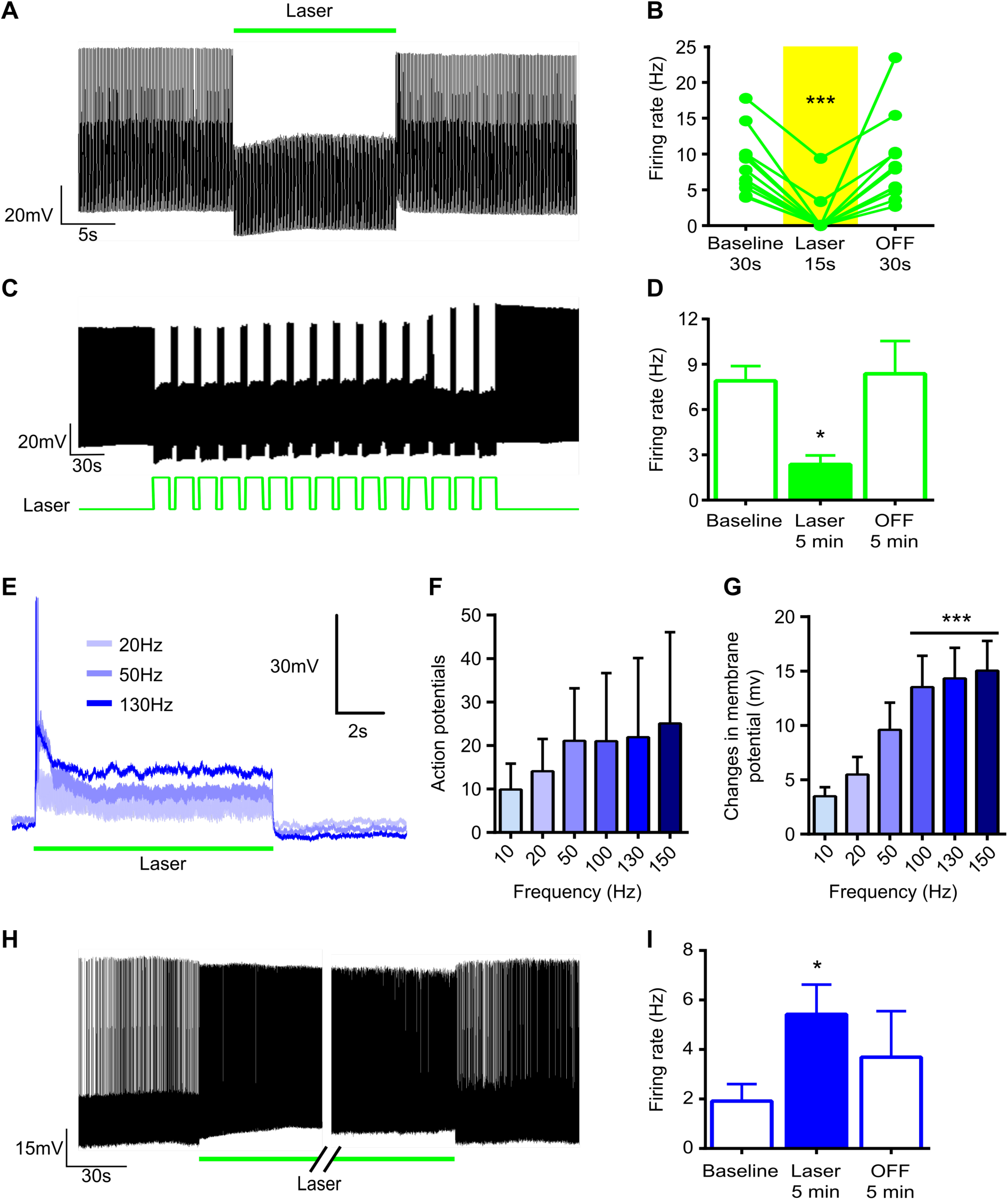
Effects of optogenetic modulation on STN neurons. **A.** Typical response recorded in a STN neuron expressing ARCHT driven at a 10Hz frequency undergoing 15s of laser inhibition. **B.** Firing rate in STN neurons expressing inhibitory opsin ARCHT was reduced by a single 15s inhibitory light pulse (Friedman test, P < 0.0001; Dunn’s post-hoc test: ***, P < 0.001, vs baseline, n = 10 cells). **C.** Example of response for a STN neuron during 5 min of laser inhibition applied with the same pattern as during behavioral experiments. **D.** Firing rate in STN neurons expressing ARCHT was reduced by 5min of application of laser pattern used during behavioral testing (Friedman test, P = 0.0207; Dunn’s post-hoc test: *, P < 0.05, vs baseline, n = 7 cells). **E.** Typical excitatory response recorded in current-clamp mode for a STN neuron expressing excitatory opsin CHETA-TC and facing 10s of laser stimulation at various frequencies: 20 (light grey), 50 (purple) and 130Hz (blue). **F.** Number of action potentials elicited during 10s of optogenetic stimulation (Friedman test, P = 0.6145, n = 10 cells). **G.** Variation of membrane potential caused by the 10s optogenetic stimulation at various frequencies (Friedman test, P < 0.0001; Dunn’s post-hoc test: ***, P < 0.001, vs 10Hz, n = 10 cells). **H.** Example of response in CHETA-TC positive STN neuron at onset (left panel) and offset (right panel) of 5min laser stimulation at 130Hz used during behavioral testing. **I.** Average changes in firing rate induced by 5 min of 130Hz laser stimulation (Friedman test, P = 0.0289; Dunn’s post-hoc test: *, P < 0.05, vs baseline, n = 8 cells).

To further assess the effect of optogenetic inhibition, we electrically stimulated STN neurons thus forcing them to emit action potentials in a stable manner (10Hz, injected current: rheobase + 20-130pA) and then applied the laser stimulation. A 15s light-pulse on STN neurons expressing ARCHT3.0 successfully inhibited neuronal firing (Fig. 5A & B). We then applied the same light pattern that was used for the behavioral experiments. The cells were thus subjected to an alternation of 15s illumination followed by 5s of obscurity during 5min (Fig. 5C). Inhibition induced by the laser reduced the firing rate of STN neurons and when the laser was stopped the cells resumed to their basal firing rate (Fig. 5C & D). This confirmed that our manipulations were sufficient to inhibit STN neurons.

The effects of optogenetic stimulation were first assessed in STN neurons with 10s of laser illumination at various frequencies (Fig. 5E). Interestingly, higher frequencies of laser stimulation did not change the number of induced action potentials compared to lower frequencies (Fig. 5F). However higher frequencies significantly increased membrane potential, suggesting this pattern increased the excitability of STN neurons (Fig. 5G). To test this hypothesis we applied 10Hz electrical stimulation (injected current: Rheobase +0-50pA) to elicit a few action potentials and we applied 130Hz laser stimulation for 5min. Firing rate of STN neurons was significantly increased by the 130Hz laser stimulation, and once the laser was turned off firing rate was no longer different from the baseline period (Fig. 5H & I). This further enforced that 130Hz optogenetic stimulation is unlikely to induce action potentials on its own, but seems to increase STN neurons excitability.

## Discussion

The data presented here show that STN optogenetic manipulations are sufficient to achieve bidirectional control on food motivation in rats. This positions the STN as a major contributor in the processes regulating sweet food reinforcement and motivation. Optogenetic inhibition mimics the effects obtained with either electric DBS or lesion of STN by increasing motivation for food and decreasing that for cocaine (Baunez et al., 2005; Rouaud et al., 2010). Optogenetic stimulation applied at 130Hz produces the mirror effect and reduces food motivation, revealing that optogenetic stimulation although applied at 130Hz results in opposite effects to those of electric 130Hz stimulation. Interestingly, STN optogenetic manipulations have transitory effect on food and cocaine intake, while electric STN DBS produces long lasting effects (Rouaud et al., 2010). This indicates that the surrounding network and afferences strongly contribute to maintain these effects, in line with previous studies showing that the hyperdirect pathway is critical in DBS treatment of some PD symptoms (Gradinaru et al., 2009; Li et al., 2012).

Because of our long lasting behavioral experiments, we used an intermittent laser modulation to lower the risk of tissue damage. Similar intensities have been used for inhibition and stimulation without any adverse consequences (Fife et al., 2017; Nieh et al., 2015). We did not observe any traces after histology that could be related to light induced damages, controls expressing EYFP were not impacted by the laser beam and animals of ‘inhibition’ and ‘stimulation’ groups were still responsive to light modulation even after several sessions, and the effect of optogenetic manipulation dissipated more or less quickly as the manipulation ceased, thus ruling out the risk of permanent alteration. Such intermittent pattern could hold a huge potential to achieve optogenetic control during prolonged experiments for future application.

We tested the influence of various parameters for optogenetic stimulation in the food experiment. Behavior was moderatly impacted by the stimulation at 5mW, and significantly reduced by the stimulation with light intensity set at 10mW. This is explained by higher depolarizing currents in STN neurons (Fig. S6F) but also because light is transmitted through a larger volume of tissue and can stimulate a larger pool of neurons (Yizhar et al., 2011).

Our electrophysiological experiments show that high frequency optogenetic stimulation increases the activity of STN neurons at cell body level. Even if optogenetic stimulation of cell bodies can lead to a different pattern once the excitation wave reaches the axon (Yizhar et al., 2011), our results advocate in favor of an effective stimulation of STN output, as it produced opposite effects to those observed after optogenetic inhibition on motivation for food. Of note, optogenetic stimulation at high frequencies of glutamatergic terminals has been used in vitro and in-vivo to induce LTD (Klavir et al., 2017), thus a similar effect could be accounted for the progressive loss of efficiency of the laser stimulation and for the rebound effect we observed during the food self-administration after several sessions of 130Hz optogenetic stimulation. In addition, our stimulation resulted in an increased activity of STN without locking it to a define rhythm, thus addressing one of the issues often raised by critics regarding optogenetic experiments, that is to determine if the activity resulting from the stimulation pattern possesses any physiological relevance (Deisseroth, 2015; Yizhar et al., 2011).

We reported increased locomotor activity consecutive to STN optogenetic inhibition (Fig. S1A). If this increased locomotor activity could explain the observed operant behavior for food reward, it is interesting to note that the laser inhibition did not change the number of responses in the inactive hole (Fig. S2C-D). Only the number of rewards and responses in the food magazine were increased by the laser inhibition (Fig. S2A-B). This suggests that even though the global activity of the animals is altered by the laser inhibition, it does not happen in an unspecific manner but remains goal-directed. Despite the fact that the first study of STN optogenetic modulation did not report any changes in locomotor activity in parkinsonian models, probably as a result of lower opsin efficiency (Gradinaru et al., 2009), more recent studies have shown that STN optogenetic inhibition could increase locomotion in PD rodent models (Yoon et al., 2016, 2014). It was also established that reducing STN glutamate transporter expression increases locomotor activity (Schweizer et al., 2014), although electric STN DBS has no effect on locomotor activity in naïve rats (Pelloux et al., 2018). Our results further extend these observations in naïve rats, and indicate that direct optogenetic inhibition of the STN is sufficient to increase locomotion. This confirms that specific inhibition of the STN leads to locomotor effect while the additional stimulation of the afferences, efferences and neighboring structures by electric DBS can counteract this effect.

Finally, opsin expression was mainly observable in the limbic and associative territories of the STN, yet optogenetic modulation affects the motor activity, this further enforces the idea that in rat territories are highly intertwined and the STN represents a site of convergence for numerous cortical projections, as suggested by former studies (Janssen et al., 2017; Kita et al., 2014).

Although STN stimulation does not alter the motivation to work for cocaine nor the locomotor activity, STN inhibition impacts both of them, in line with previous studies (Baunez et al., 2005; Rouaud et al., 2010). Two elements can explain this finding. First, our optogenetic stimulation seems to increase STN excitability, and that increase might be insufficient to affect cocaine related motivation and locomotion. Second, the STN seems highly sensitive to specific ranges of oscillations. For instance in the context of PD, DBS at low frequency (< 20Hz) has been reported to worsen the symptoms, while high frequency DBS exerts therapeutic effects (Eusebio et al., 2008; Timmermann et al., 2004). Very similarly, in the STN, very low frequencies (1-4Hz) have been closely tied to the severity of obsessive compulsive disorders (Welter et al., 2011). Finally, recent data from our group indicate that low frequency oscillations (Theta 8-12Hz and Beta 13-30Hz) develop in the STN during extended cocaine use, and that blocking these oscillations with DBS or lesion prevents escalation and re-escalation of cocaine intake (Pelloux et al., 2018). Thus, the key point to affect cocaine motivation and motor activity through STN stimulation probably lies within very specific ranges of oscillations or a specific driving rhythm rather than a general increase of activity.

Overall the present work emphasizes some of the specific contributions of STN in reward related processes: food and cocaine self-administration, as well as locomotor behavior. By providing data on three different behaviors that can be deeply impacted by DBS, we hope to offer a large comparative framework for the study of DBS effects. Since DBS is progressively applied for an increasing number of psychiatric pathologies such as depression, obsessive-compulsive disorders and addiction (Hamani et al., 2017; Mallet et al., 2008; Pelloux and Baunez, 2013), identifying the contribution of the structure itself and that of the networks supporting therapeutic effects appears critical to reduce the risks of side effects and refine the parameters and strategies used to target brain structures.

## Material and Methods

### Animals

Lister hooded male rats, 380-400g, were used in this study. They were housed in pairs; maintained in an inverted 12h dark/light cycle (light onset at 7 p.m.) in temperature-controlled room, with unlimited access to water. All experiments were carried out during the dark phase. For the food-rewarded experiments, rats were kept under moderate food restriction (15-16g/rat, 75-80% of normal daily intake). For the other experiments animals had no food restriction. All animal care and use conformed to the French regulation (Decree 2010-118) and were approved by the local ethic committee and the University of Aix-Marseille (saisine #3129.01).

### Viral vectors

We used AAV5 (UNC Vector core, Chapel Hill, USA) to transfect STN neurons, allowing the expression of recombinant protein under the CamKIIa promoter with the following constructs: AAV5-CaMKII-hChR2 (E123T/T159C)-p2A-EYFP-WPRE (CHETATC) for stimulation groups, AAV5-CaMKII-ArchT3.0-p2A-EYFP-WPRE for inhibition groups and AAV5-CaMKII-EYFP for the control groups.

### Surgery

Animals were anesthetized with a combination of Ketamine (Imalgen, Merial, 100mg/kg, s.c.) and medetomidine (Domitor, Janssen, 30mg/kg, s.c.).

#### Optogenetic surgery

Animals were placed in a Kopf stereotaxic apparatus to receive a bilateral 0.5 μL injection of virus (0.16μL/min) using injectors placed at the following coordinates relative to bregma: AP: -3.7 mm; ML: ±2.4 mm; DV: -8.4 mm (from skull) according to the Paxinos and Watson atlas (Paxinos and Watson, 2005). Optic fibers were implanted 0.5mm above each injection site and secured within a head-cap made of dental cement on the skull.

#### Catheter surgery

A homemade silicone catheter was inserted 2.5-3.0cm in the right jugular vein and secured with ligatures and exited dorsally between the scapulae through a guide cannula secured with dental cement. Blood reflux in the tubing was checked to confirm correct placement of the catheter. Catheters were also daily flushed with heparin (Sanofi, 3g/L) and antibiotic enroflorilexine (Baytril, Bayer, 8g/L) in 0.9% saline to prevent blood clots and infection during the recovery period, and were also flushed before and after each self-administration session until the end of experiments.

After surgery, all rats were awakened with an injection of atipamezol (Antisedan, Janssen, 0.15mg/kg, *i*.*m*.). Preventive long-acting amoxicillin treatment was applied (Duphamox LA, Pfizer, 100mg/kg, *s*.*c*.). They were allowed to recover for at least 7 days, during which they were daily monitored. To allow for satisfying level of expression of the opsins, a waiting period of 3 weeks was applied before any light delivery.

### In-vivo light delivery

Implants were built with 230 μm optic fibers (NA 0.22, Thorlabs) glued to 2.5mm ceramic ferrules (Thorlabs) using epoxy. Ferrules were slightly grinded with a Dremel to improve the contact with the dental cement. Light stimulation was performed during experiments using 200mW 532nm DPSS laser controlled with a signal generator. Optic rotary joints (DoricLenses) relieved the torsion due to animal movements and bilateral stimulation was achieved using an optic coupler (Thorlabs, FCMM625-50A). Light output was assessed before each experiment with a power meter (Thorlabs, PM20A) and parameters for stimulation were: 10mW, 2ms pulse, 130Hz; and for inhibition: 5mW, 15s pulse, 5s interval between pulses (resulting in a 0.2Hz pattern), unless specified otherwise.

### Food self-administration

Rats were first trained during10-15 days to perform the fixed-ratio 1 for sucrose pellets in standard rat operant chambers (MedAssociates) with an active hole paired with the delivery of the reward and illumination of an associated cue light and an inactive hole, whose activation had no programmed consequences. The ratio was progressively increased until fixed ratio 5 (FR5), stability of performance was evaluated during 3 days before incrementing to the next ratio. Sessions of FR5 lasted 30 min. In the case of progressive ratio (PR) experiments, the ratios followed an arithmetically increasing schedule in steps of five, with three repetitions of each step (i.e., 1, 1, 1, 5, 5, 5, 10, 10, 10…). The sessions ended after 90min or after 5min without any response in the active hole and the last ratio completed was defined as the animal ‘breakpoint’.

### Cocaine Self-Administration

The intravenous cocaine self-administration was conducted in the same operant boxes described above; the stainless steel guide cannula of the catheter was connected through steel-protected Tygon tubing to a swivel (Plastics One) and then a syringe filled with cocaine placed on an infusion pump (MedAssociates). Self-administration experiments used a dose of 250μg/90μL per cocaine injection (Coopérative pharmaceutique française). The daily sessions lasted for 2 h for FR5 or maximum 4 h for progressive ratio. In PR sessions, to limit the number of injections, the ratios for cocaine reward followed the modified equation of Roberts 1, 3, 6, 10, 15, 20, 25, 32, 40, 50, 62, 77, 95, 118, 145, 178, 219, etc. (Depoortere et al., 1993). If the rats failed to complete a ratio within 60 min, the session ended.

### Spontaneous‐ and cocaine-induced locomotor activity

Locomotor activity was measured as the distance traveled (in m) in a circular homemade Perspex open-field (60 cm diameter) with a video tracking performed with Bonsai (Open Ephys). Details of the procedure can be found in the supplementary material.

### Electrophysiology

Full details regarding solutions and preparation of adult brain slices can be found in the supplementary materials and were adapted from other studies (McDevitt et al., 2014; Ting et al., 2014). Adult rats were anesthetized and perfused with ice-cold artificial cerebrospinal fluid. 200 μm coronal slices containing the STN were prepared with a vibratome (1200S, Leica). Cells were visualized on an upright microscope with infrared differential interference contrast and fluorescence microscopy (Olympus, BX51WI). Recordings were made with pClamp 10.3 software using a MultiClamp 700B amplifier set with 4 kHz low-pass Bessel filter and 10 kHz digitization (Molecular Devices, Sunnydale, CA). Laser modulation was applied using a 532nm DPSS laser. The light beam was directed at the preparation using an optic fiber submerged into the bath. Laser intensities were adjuted to match those used in behavioral experiments.

To assess the influence of laser modulation on firing pattern, cells were imposed a given rhythm through current injection. Cells expressing inhibitory opsins ARCHT3.0 were driven at 10Hz, with injected current intensity corresponding to rheobase + 20-130pA in order to induce stable and robust firing pattern. Cells expressing excitatory opsins CHETATC were driven at 10Hz, intensity set at +0-50pA to elicit a few action potentials in a stable pattern. Stability of the excitation was assessed trough a 2min baseline recording, then we applied the same laser pattern used for behavioral experiments for 5min and the cells were monitored for another 5min with the laser turned OFF.

Extended details for histological controls and statistic analyses can be found in the supplementary materials.

## Acknowledgments

The authors thank Drs. F Brocard and R Bos for allowing use of their Patch clamp equipment and animal facility personal for technical support.

## Supplemental information

### Electrophysiology

Animals were anesthetized and perfused intra-cardiacally with ice-cold artificial cerebrospinal fluid (ACSF). 200 μm coronal slices containing the STN were prepared in ice-cold ACSF. After being cut, the slices were maintained for 10 minutes at 33°C and then transferred to holding ACSF at room temperature. ACSF used for perfusion, cutting, and recovery contained NMDG as a sodium substitute and contained, in mM: 92 NMDG, 2.5 KCl, 1.25 NaH2PO4, 30 NaHCO3, 20 HEPES, 25 glucose, 2 thiourea, 5 Na-ascorbate, 3 Na-pyruvate, 0.5 CaCl2 and 10 MgSO4-7H2O (pH: 7.35). ACSF used for holding slices prior to recording was identical, but contained 92 mM NaCl instead of NMDG and contained 1 mM MgCl and 2 mM CaCl2. ACSF used to perfuse slices during recording was maintained at 31°C and contained, in mM, 125 NaCl, 2.5 KCl, 1.25 NaHPO4, 1 MgCl, 2.4 CaCl, 26 NaHCO3 and 11 glucose. All ACSF preparations were saturated with 95% O2 and 5% CO2.

Cells were patched using glass pipets with resistance 3.5-5.0MΩ, filled with internal solution containing, in mM, 140 potassium gluconate, 5 NaCl, 2 MgCl2, 10 HEPES, 0.5 EGTA, 2 ATP, 0.4 GTP (pH: 7.35). Series resistance was monitored during experiment with 10mV hyperpolarizing pulses and maintained below 20MΩ, or cells were discarded. Firing pattern of STN neurons was determined by injecting depolarizing currents during 500ms with 20pA increment steps.

Cells were optically stimulated through a 200μm fiber placed in the bath with light beam aimed at the STN. Intensity was set at 10mW for stimulation and 5mW for inhibition experiment, except when the influence of the light intensity was assessed. For experiments testing the influence of light intensity and pulse width, cells were held at -60mV in current-clamp configuration and discrete light pulses were applied. When testing the influence of light intensity, pulse duration was set to 15s and 1s for inhibition and stimulation groups, respectively. Testing the influence of the pulse duration was performed at constant light intensity: 10mW (Stimulation) and 5mW (Inhibition).

### Spontaneous‐ and cocaine-induced locomotor activity

Locomotor activity was measured as the distance traveled (in m) in a circular homemade Perspex open-field (60 cm diameter) with a video tracking performed with Bonsai (Open Ephys), recorded on a PC and analyzed offline. Rats were placed in the open-field for 30 min of habituation and were connected to the optic fiber cable and further recorded for 30 min during which they underwent laser stimulation for 2 bins of 5 min each. Their spontaneous locomotion was measured. For the measure of cocaine-induced locomotion, the animals were injected at the end of the habituation with either 0.9% NaCl (1 ml/kg) or one of the various doses of cocaine (5; 7.5 and 10 mg/kg, *s.c*.) whose order was counterbalanced in a pseudo-latin square manner, with a five-day interval between two injections to allow for washout. The doses were chosen according to previous studies in line with their reinforcing effect associated with limited effect on locomotion to avoid stereotyped behavior in case of potentiation by the STN manipulation (Baunez et al., 2005).

### Histology

At the end of the experiment, the rats were deeply anesthetized with pentobarbital (Dolethal, *i.p.*) and perfused intracardiacally with 4% paraformaldehyde dissolved in PBS. The brains were extracted and after cryo-protection in 30% sucrose, they were frozen into liquid isopentane (−80°C) to be further cut in 40μm coronal slices with a cryostat

For the optogenetic experiments, the brain sections were examined for optic fibers location and for native fluorescence expression with an epifluorescence microscope (Zeiss, Imager.z2) immediately after being cut, except for the CHETA-TC animals, in which native fluorescence was present but required immuno-staining to control the exact boundaries of the viral expression.

Sections underwent a 90 min permeation step (PBS, 1% bovine serum albumin (BSA) 2% normal goat serum (NGS), 0.4% TritonX-100), 3×5min PBS washes, incubation with primary antibody (mouse anti-GFP, A11120, Life technologies; 1:200, in PBS 1% BSA, 2% NGS, 0.2% TritonX-100) at 4°C overnight. The sections were then washed 3×5min with PBS followed by 2h incubation at room temperature with secondary antibody (Goat anti-mouse Alexa 488, A11011, Life technologies, 1:400 in PBS 1% BSA, 2% NGS), and finally washed 3×5min with PBS before being mounted on to glass slides with homemade mounting medium.

67 animals were used in the optogenetic procedure, among which 2 were excluded after loss of optic fiber implants, 3 for absence of fluorescence in either one or both STN, and 1 due to optic fiber misplacement.

### Statistical analyses

Data are expressed as mean ± SEM with the sample size indicated for each group. Analyses were performed with Prism (GraphPad) and Matlab (Mathworks) software. Behavioral data were analyzed with mixed-model design ANOVA (mixed ANOVA) with groups as between factor and sessions, time bins, or light parameters as a repeated within factor. Follow-up analysis was conducted using Sidak’s post-hoc test for multiple comparisons or Fisher’s post-hoc test for single comparison. Data regarding cocaine-induced locomotion were analyzed with 2-way Repeated Measure ANOVA with dose and time bins as repeated within factors. Two samples comparisons were analyzed with non-parametric Mann-Whitney test. Electrophysiological data were analyzed with Friedman test followed by Dunn’s post-hoc test, variations in cells properties were analyzed with Kruskall-Wallis test or mixed ANOVA. Only *P*-values < 0.05 were considered significant.

---

**Supplementary Fig. 1:**
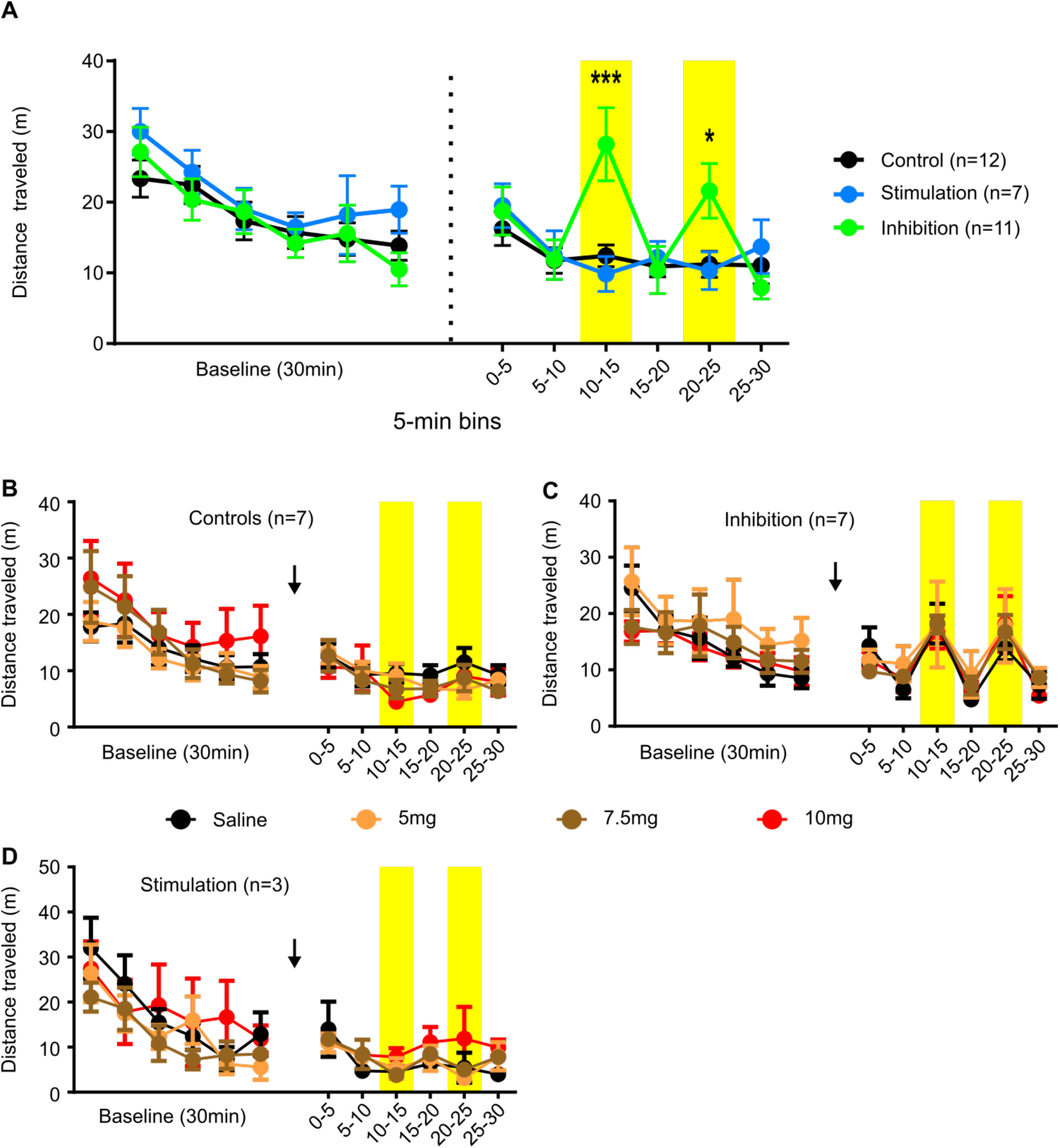
Effect of STN optogenetic modulation on rat spontaneous and cocaine-induced locomotion. **A.** Locomotor activity expressed in distance travelled (in m) per 5 min bins during the 30min of habituation (left) and 30min of test (right) with 2 bins of laser activation (yellow) for the control (black dots), stimulation (blue dots) and inhibition (green dots) groups. There was a significant interaction effect when comparing ‘inhibition’ group to ‘control’ (mixed ANOVA, F Time bins x subjects (5, 105) = 8.576, P < 0.0001; Sidak’s post-hoc test: * p < 0.05, *** p < 0.001) while no effect was observable when comparing ‘stimulation’ and ‘control’ groups (mixed ANOVA, F _Time bins x subjects_ (5, 85) = 1.253 P = 0.2920). **B – D.** Effect of STN optogenetic modulation on cocaine-induced locomotion tested with various doses of cocaine: 5mg/kg (orange), 7.5mg/kg (brown), 10mg/kg (red) and compared to the saline control (black) in control (**B.**), inhibition (**C.)** and stimulation (**D.)** groups. There was no effect of the cocaine dose on the locomotor activity in any of the 3 groups: ‘control’, ‘inhibition’ and ‘stimulation’ (Repeated Measure 2-way ANOVA, F _Time_ bins x dose (15, 120) = 1.288, P = 0.2202; F Time bins x dose (15, 120) = 0.4931, P = 0.9402; F _Time_ bins x dose (15, 40) = 0.7500, P = 0.7207, respectively). The laser beam activation produced an increase in locomotor activity in the ‘inhibition’ group regardless of the dose of cocaine (Repeated Measure 2-way ANOVA, F laser bins (5, 120) = 17.26 P < 0.0001).

**Supplementary Fig. 2:**
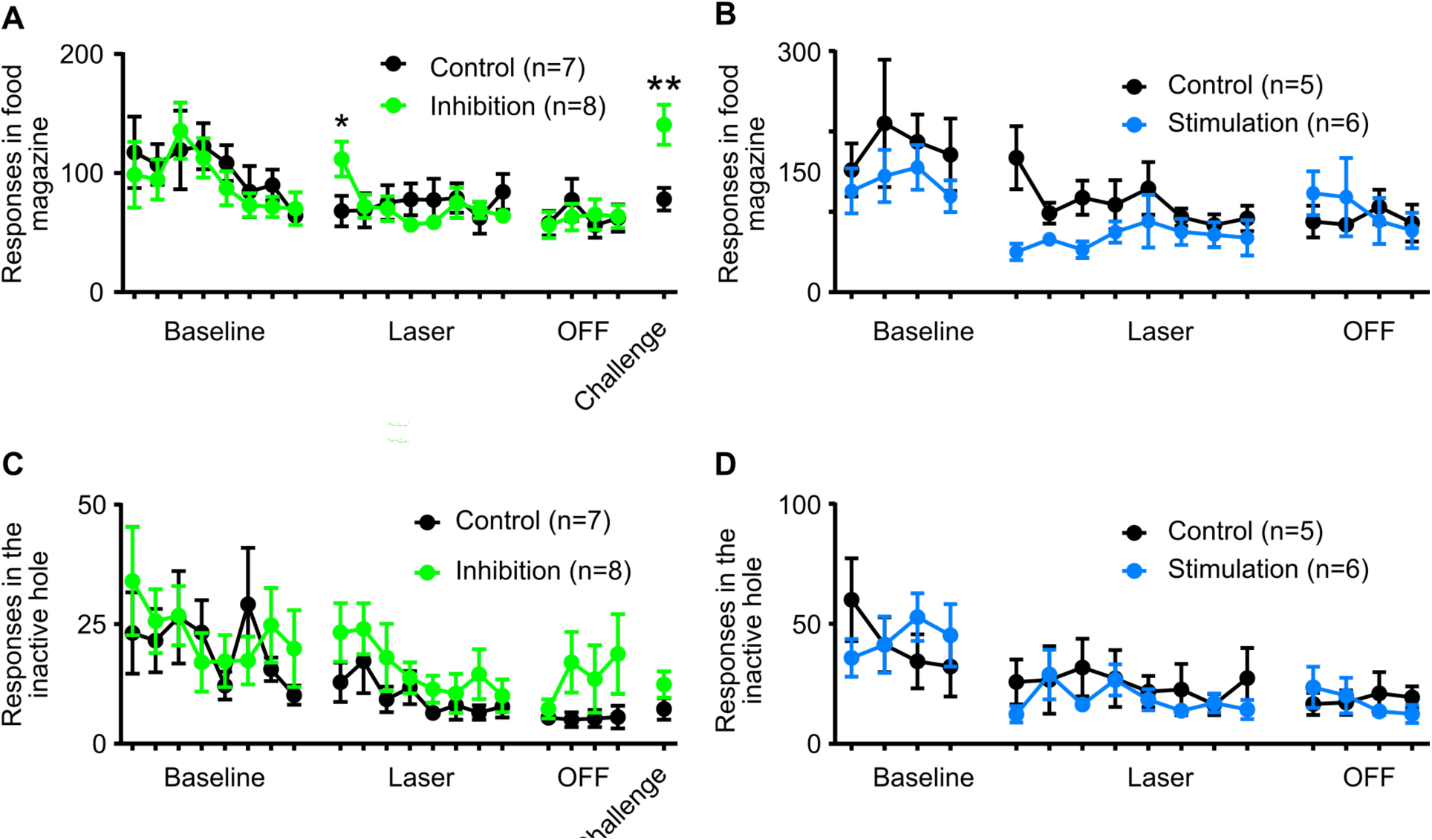
Effects of STN optogenetic modulation on food magazine visits and inactive hole responses recorded in the progressive ratio (PR) task illustrated for each session (8 sessions of baseline, 8 sessions with laser ON (i.e. Laser), 4 sessions with laser OFF (i.e. OFF) and 1 session with laser ON again (i.e. challenge)). **A.** Influence of STN optogenetic inhibition on the number of responses made in the food magazine. When the laser was turned on, there was a significant increase in food magazine responses at baseline-laser transition (1^st^ session of the laser ON period)(mixed ANOVA, F sessions x sub_jects_ (1, 13) = 8.809, P = 0.0109) and at off-challenge transition (mixed ANOVA, F sessions x subjects (1, 13) = 7.968, P = 0.0144). **B.** Influence of STN 130Hz optogenetic stimulation on the responses made in the food magazine. The 130 Hz stimulation induced a strong trend towards decreased number of responses in the stimulated animals (mixed ANOVA, F _Group_ (1, 9) = 4.864, P = 0.0549). **C.** Effect of STN optogenetic inhibition on the number of responses made in the inactive hole. The number of erroneous responses remained unaffected either during baseline-laser transition (mixed ANOVA, F sessions _x_ subjects (1, 13) = 0.008801, P = 0.9267) or during the laser challenge (mixed ANOVA, F sessions x subjects (1, 13) = 1.267, P = 0.2807). **D.** Effect of STN optogenetic 130 Hz stimulation on the number of responses made in the inactive hole. The number of erroneous responses was not affected by the 130Hz optogenetic stimulation (mixed ANOVA, F sessions _x_ subjects (1, 9) = 3.286, P = 0.1033). Fisher’s post-hoc test: * p < 0.05, ** p < 0.01, vs control.

**Supplementary Fig. 3:**
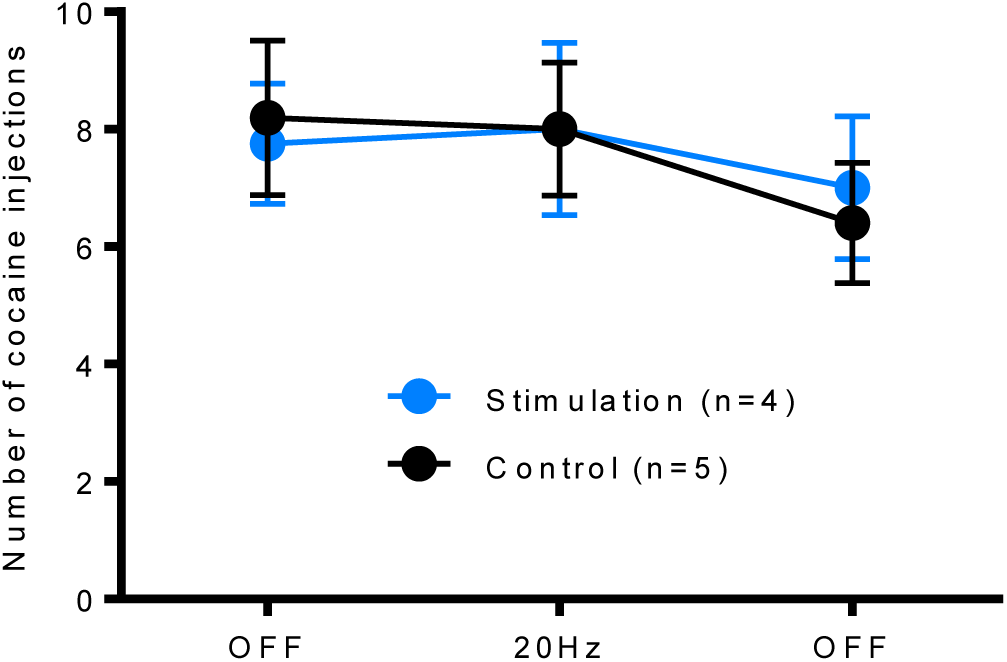
Effect of acute 20 Hz optogenetic stimulation applied during one session of progressive ratio for cocaine. The motivation of the animals (controls in black and stimulated in blue) is expressed in number of injections obtained during one session with laser OFF, one session with laser ON (20 Hz) and one last session with laser OFF. The acute 20Hz optogenetic stimulation did not alter motivation for cocaine (mixed ANOVA, F sessions x subjects (2, 14) = 0.3486, P = 0.7116).

**Supplementary Fig. 4:**
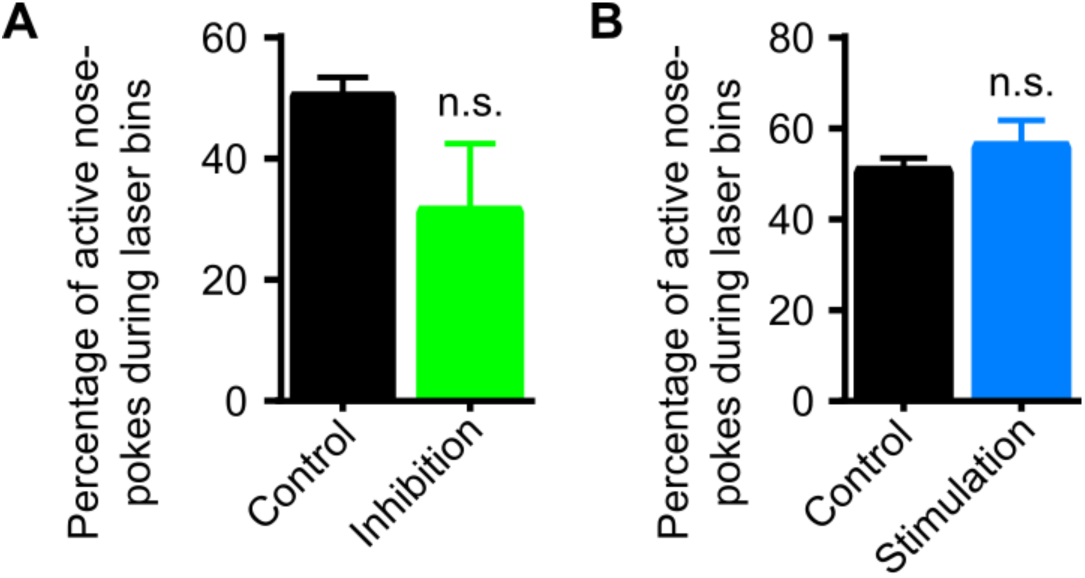
Evaluation of the percentage of active nose-pokes while the laser was activated. **A.** The percentage of active nose-poke when the laser beam was turned ON (i.e. during laser bins) was not modified compared to controls (Mann Whitney test, P = 0.2857). **B.** Laser stimulation did not alter the pattern of nose-poke of stimulated animals compared to the control group (Mann Whitney test, P = 0.4127)

**Supplementary Fig. 5:**
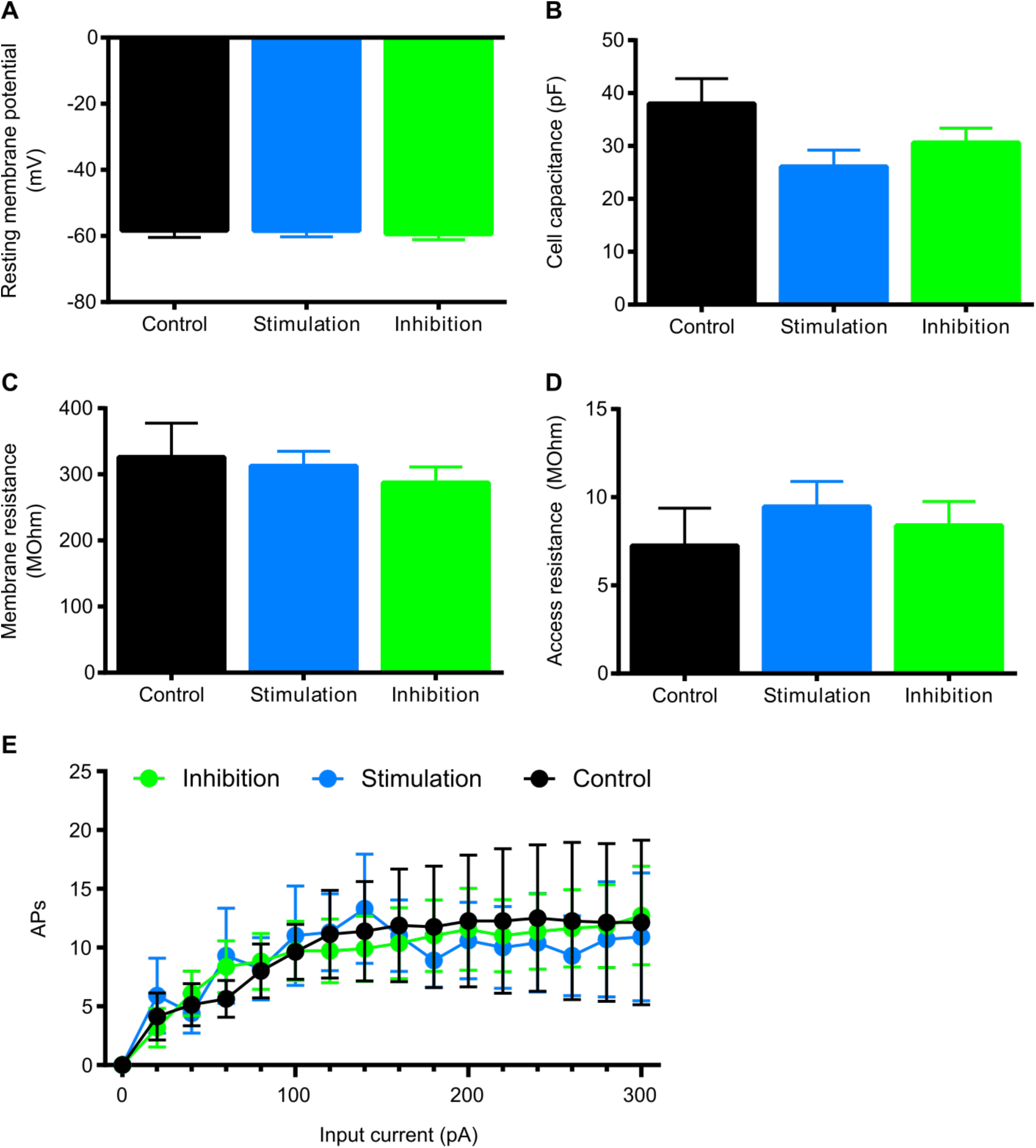
Cells properties of STN neurons after viral infection recorded in whole cell configuration for controls (EYFP, n=8 cells, black bars), stimulation (CHETA-TC, n=10 cells, blue bars) and inhibition (ARCHT, n=11 cells, green bars). Cell resting membrane potential (**A.,** Kruskal-Wallis test, P = 0.8214), cell capacitance (**B.,** Kruskal-Wallis test, P = 0.0644), membrane resistance (**C.,** Kruskal-Wallis test, P = 0.7397), and access resistance (**D.,** Kruskal-Wallis test, P = 0.3158) were unaffected. **E.** The average number of action potentials (APs) elicited by a 500ms step of hyperpolarization was not modified due to the presence of the opsins (**E.**, mixed ANOVA, F Frequency x Group (30, 390) = 0.2480, P > 0.9999).

**Supplementary Fig. 6:**
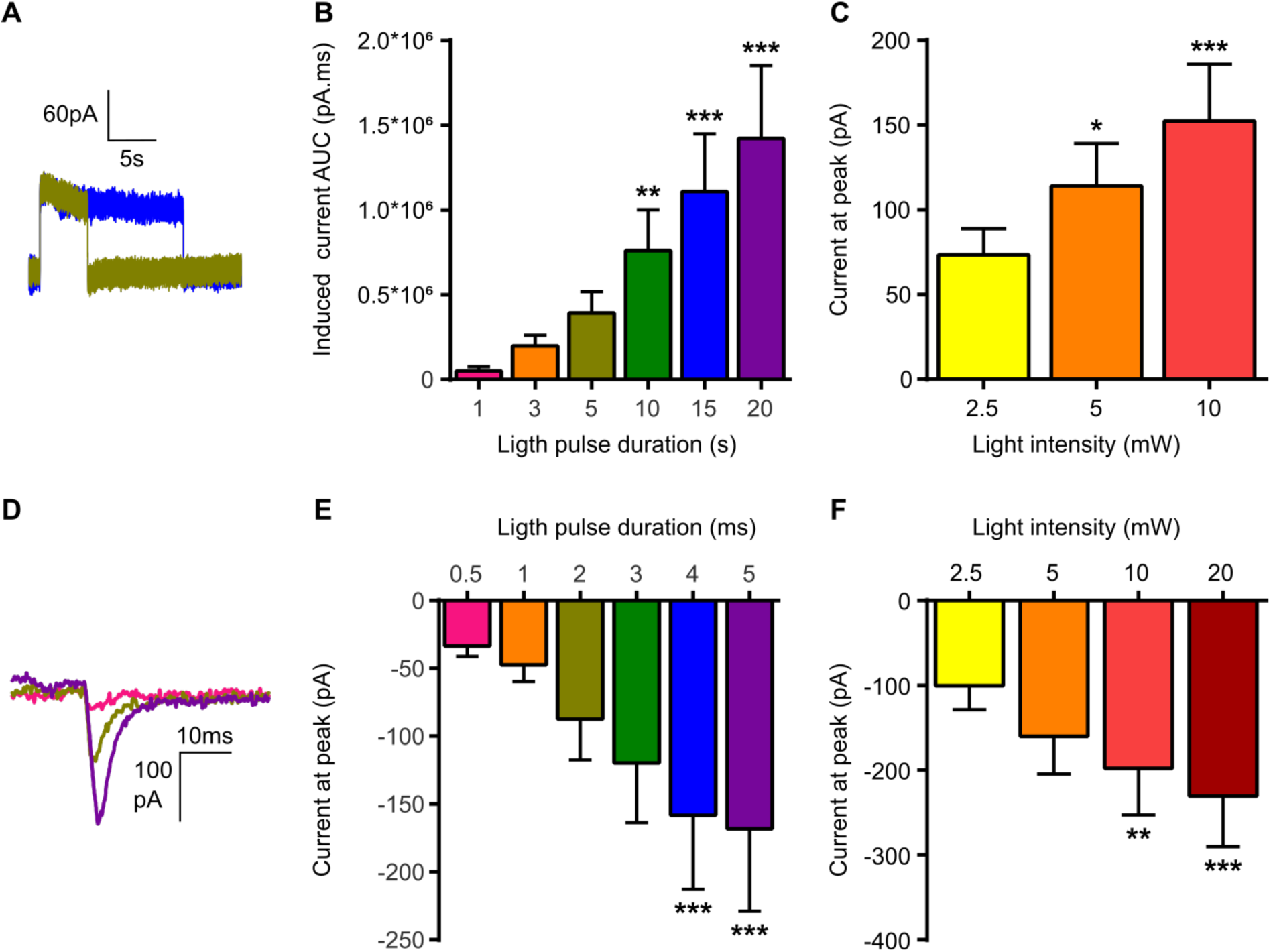
Evaluation of light stimulation parameters in cells expressing inhibitory opsins (ARCHT, n=11 cells) or excitatory opsins (CHETA-TC, n=10 cells). **A.** Example of light induced inhibitory currents in one STN neuron expressing ARCHT by a 5s (brown) and 15s (blue) pulse. **B.** Average area under the curve (AUC) calculated for different durations of light pulses with intensity set at 5mW (Friedman test, χ² (5) = 51.88, p < 0.0001, Dunn’s post-hoc test: **, p < 0.01; ***, p < 0.001; vs 1s pulse). **C.** Average current induced by a single 15s pulse applied with different light intensities (Friedman test, χ² (2) = 22.00, p < 0.0001; Dunn’s post-hoc test: *, p < 0.05; ***, p < 0.001). **D.** Example of induced depolarizing current in CHETA-TC cells induced by a 0.5ms (pink), 2ms (brown) or 5ms (purple) light pulse. **E.** Average induced current in CHETA-TC cells depending on the duration of the light pulse with intensity set at 10mW (Friedman test, χ² (5) = 46.00, p < 0.0001; Dunn’s post-hoc test: ***, p < 0.001; vs 0.5ms pulse). **F.** Induced current by a 1s light pulse as a function of light intensity (Friedman test, χ² (3) = 28.92, p < 0.0001; Dunn’s post-hoc test: **, p < 0.01; ***, p < 0.001; vs 2.5mW)

